## Supplementary figures and images for "BUB-1 and CENP-C recruit PLK-1 to Control Chromosome Alignment and Segregation During Meiosis I in *C. elegans* Oocytes"

### Figure 1 - Figure supplement 1

#### Figure 1-figure supplement 1

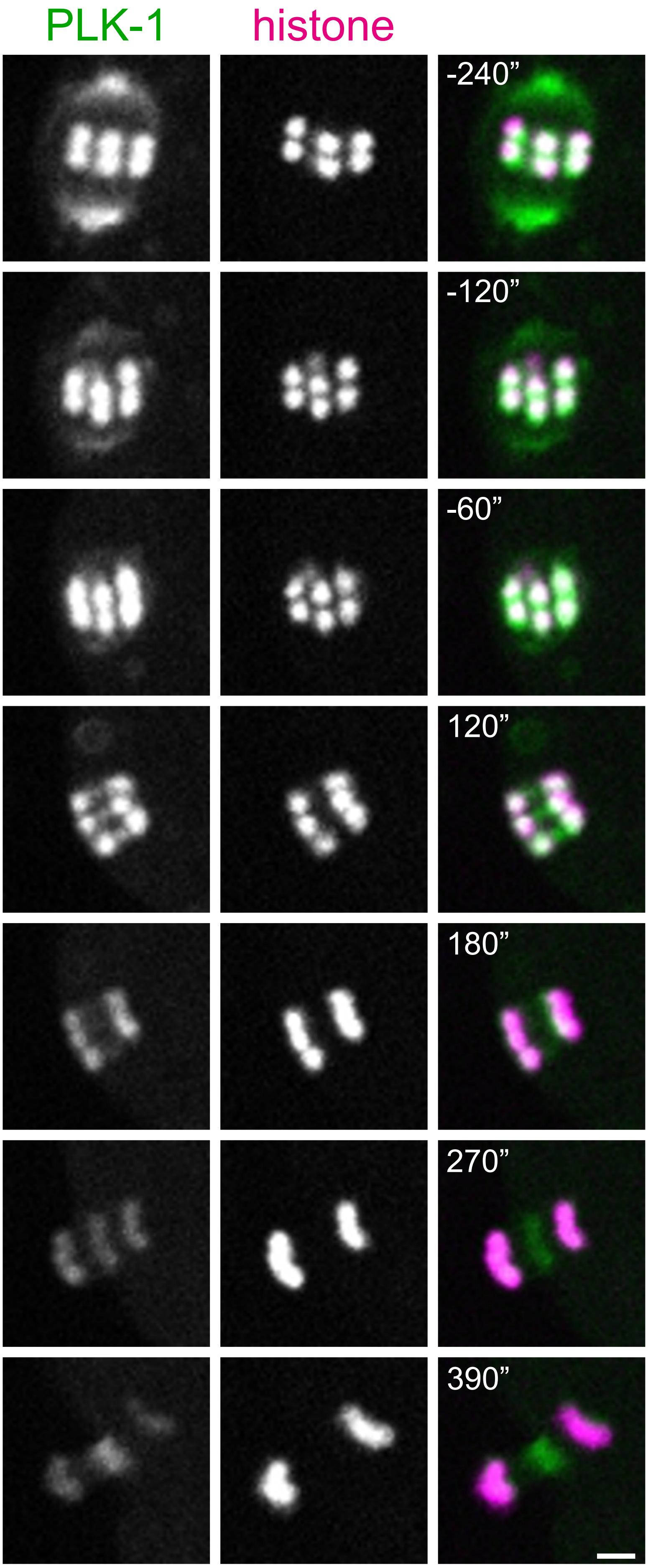


PLK-1::sfGFP; mCherry::histone

### Figure 1 - Figure supplement 2

#### Figure 1-figure supplement 2

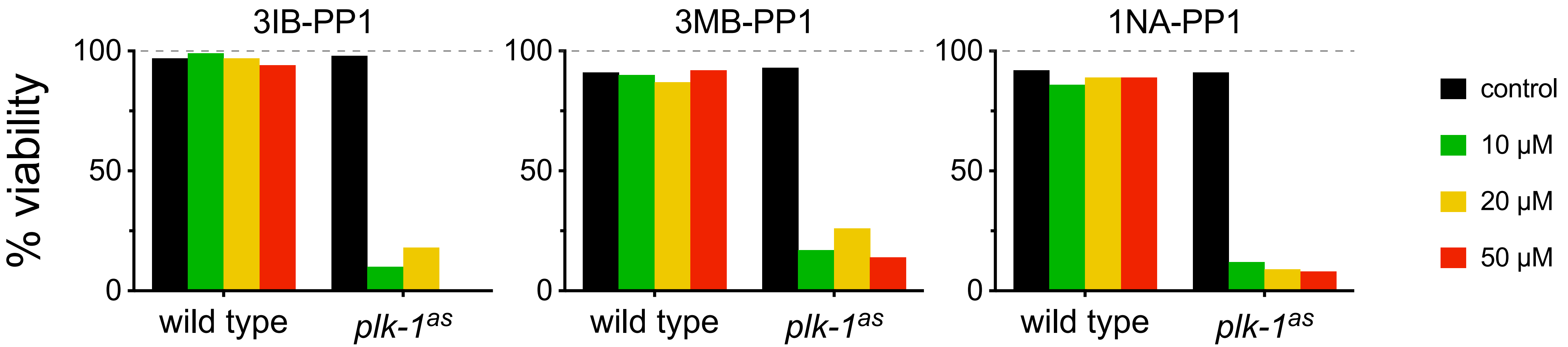

### Figure 1 - Figure supplement 3

#### Figure 1-figure supplement 3

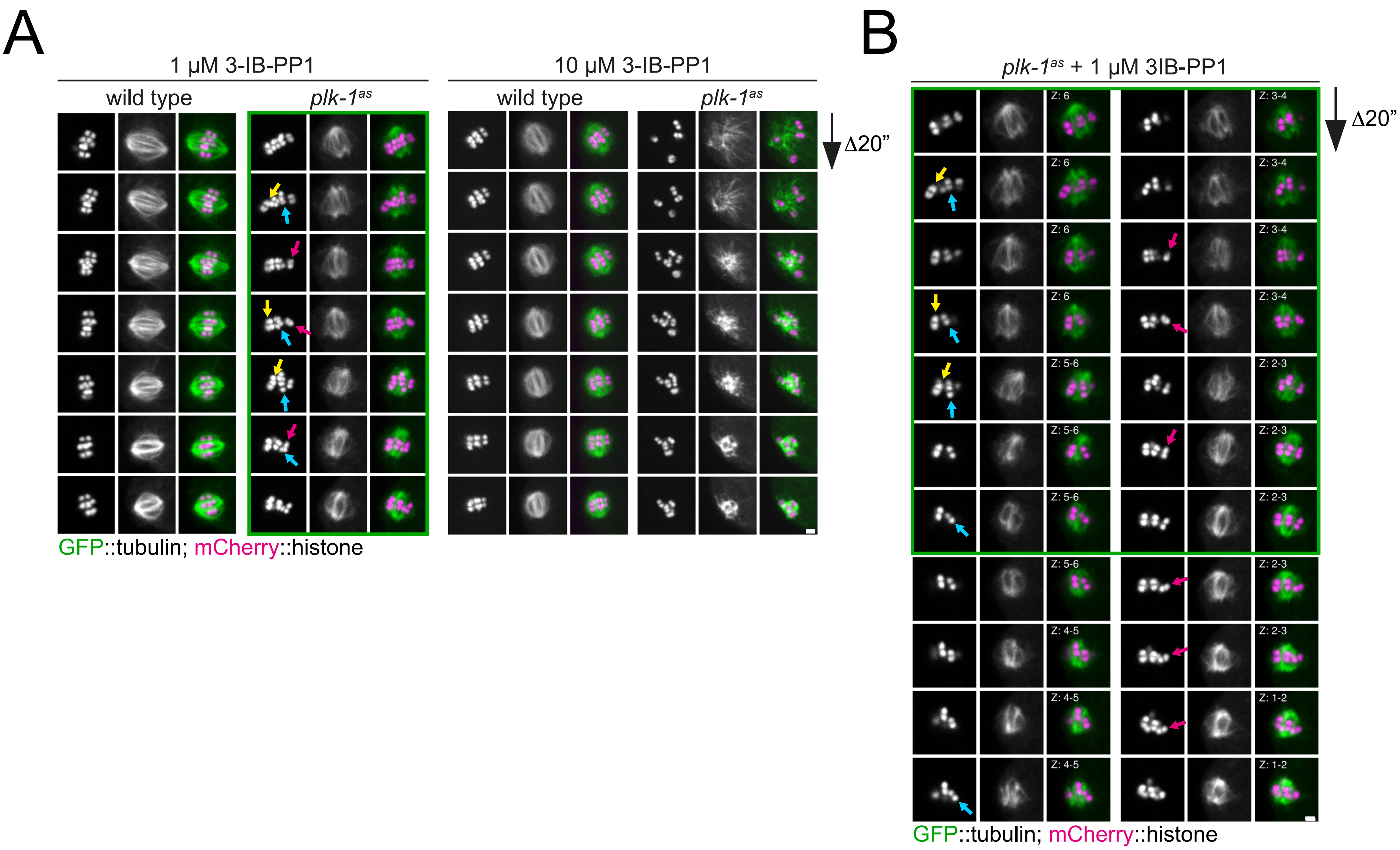

### Figure 2 - Figure supplement 1

#### Figure 2-figure supplement 1

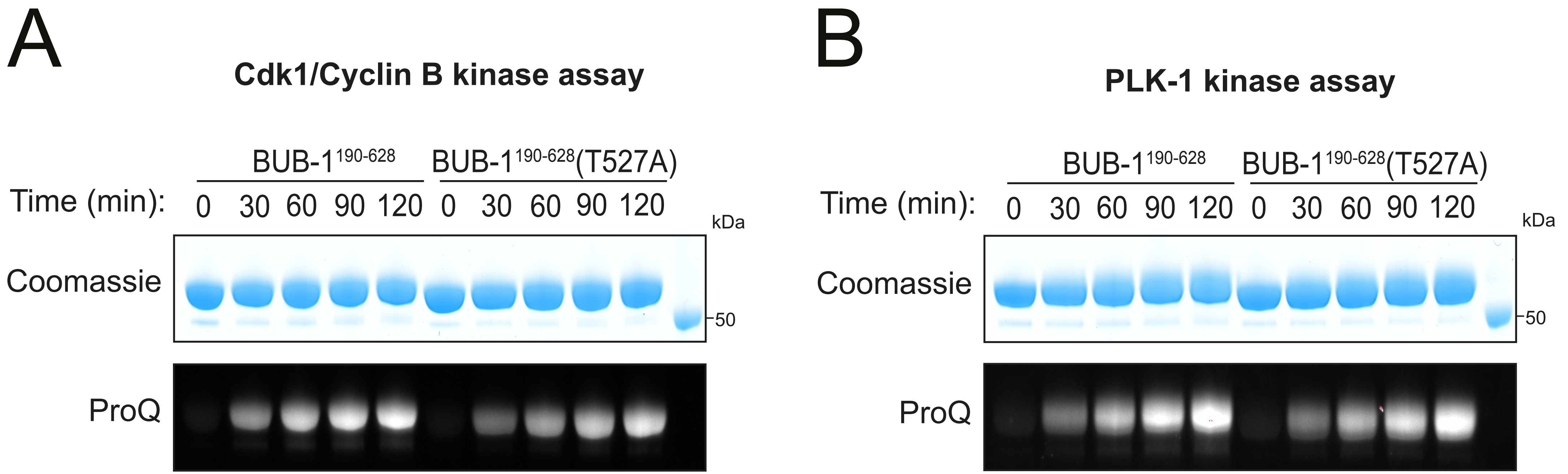

### Figure 5 - Figure supplement 1

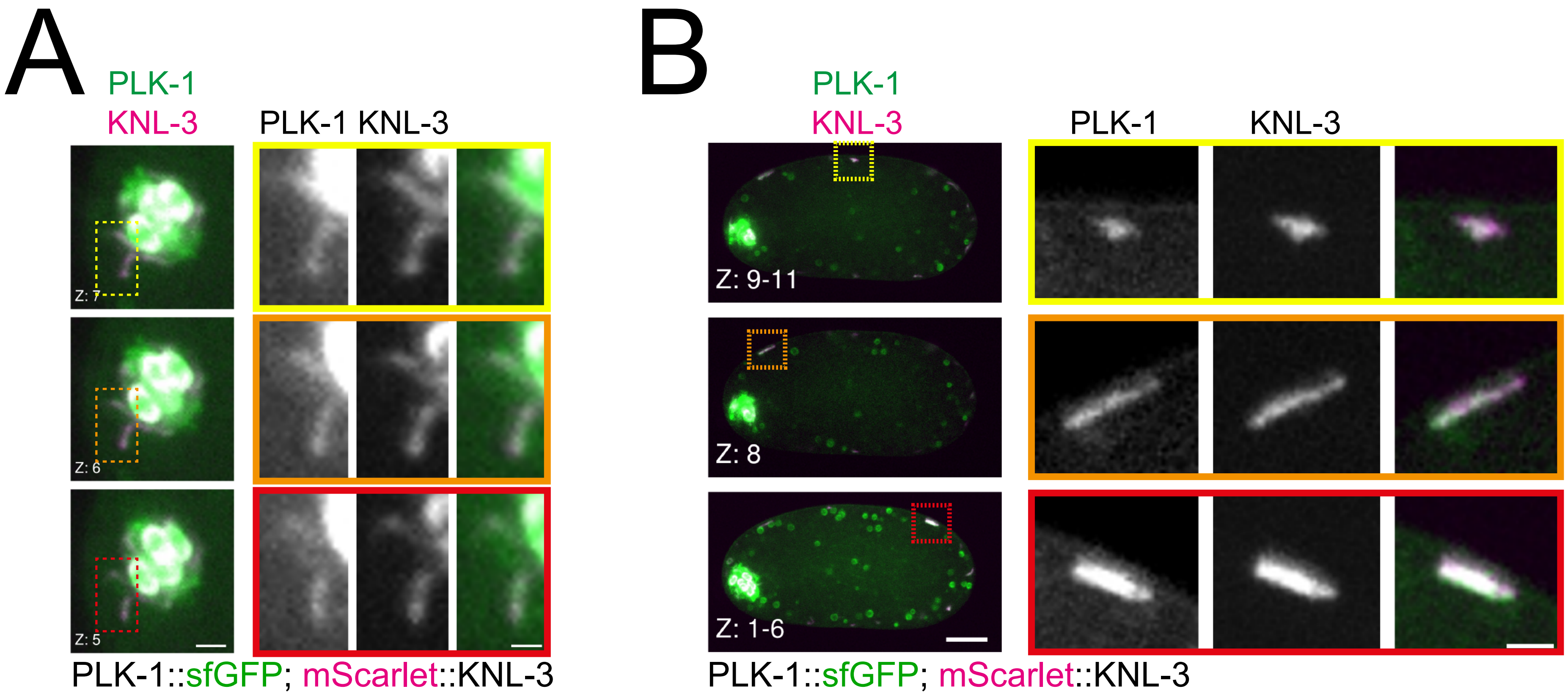

### Figure 8 - Figure supplement 1

#### Figure 8-figure supplement 1

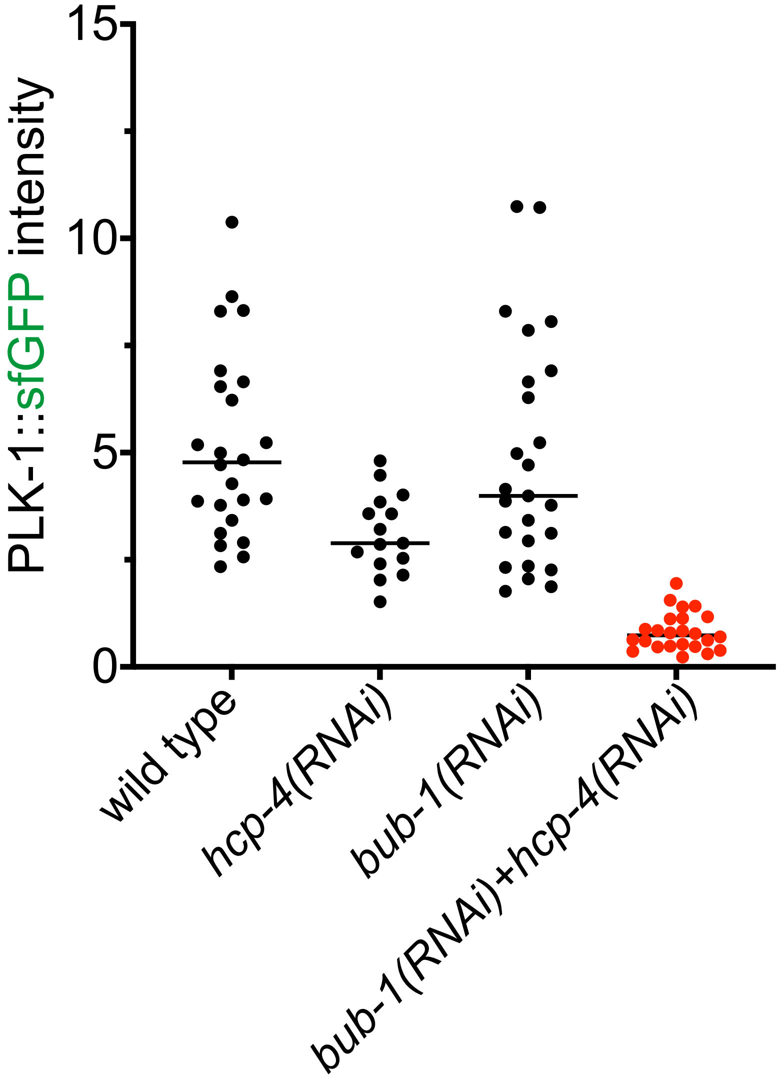

### Figure 8 - Figure supplement 2

#### Figure 8-figure supplement 2

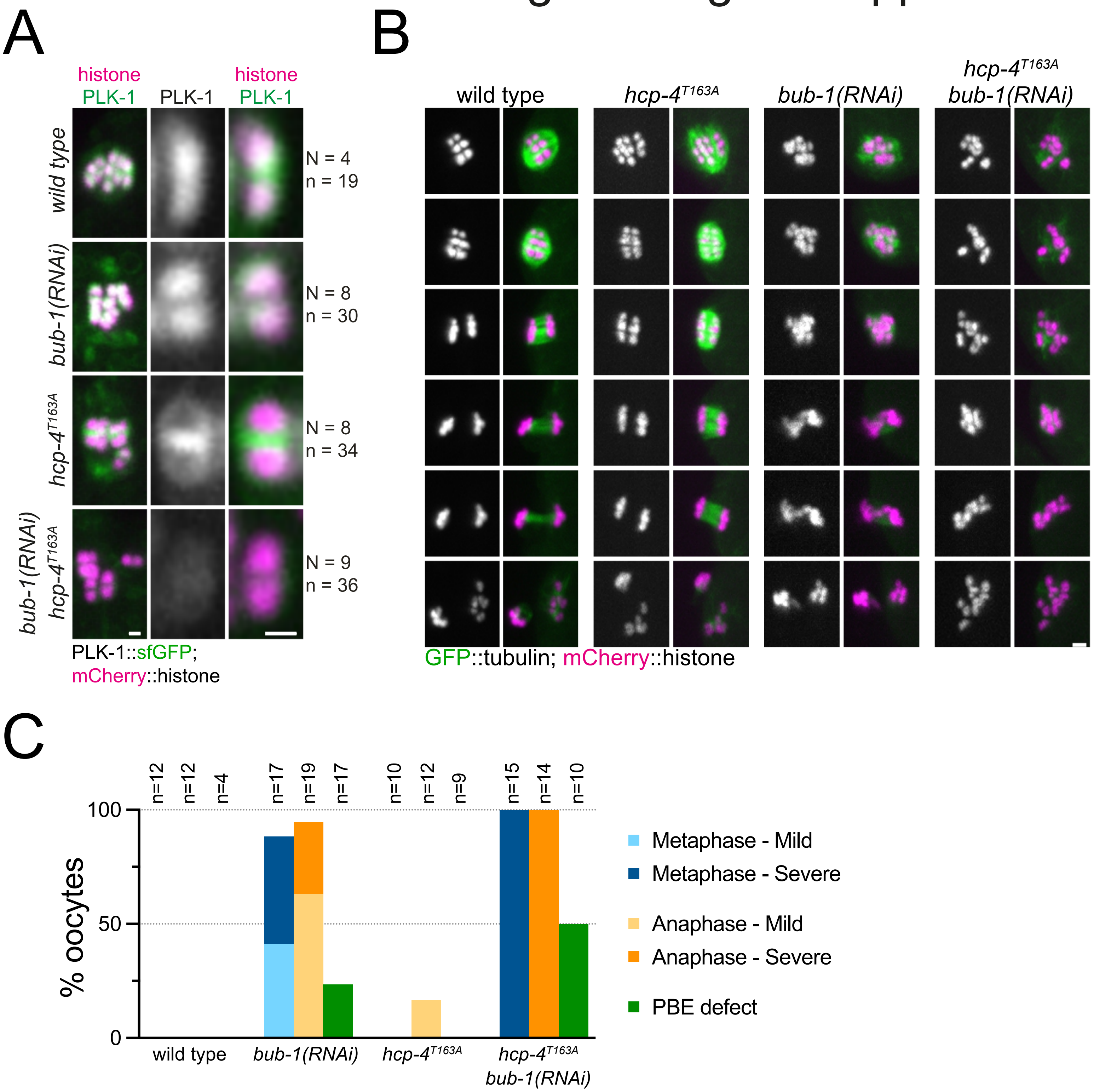

### Figure 9 - Figure supplement 1

#### Figure 9-figure supplement 1

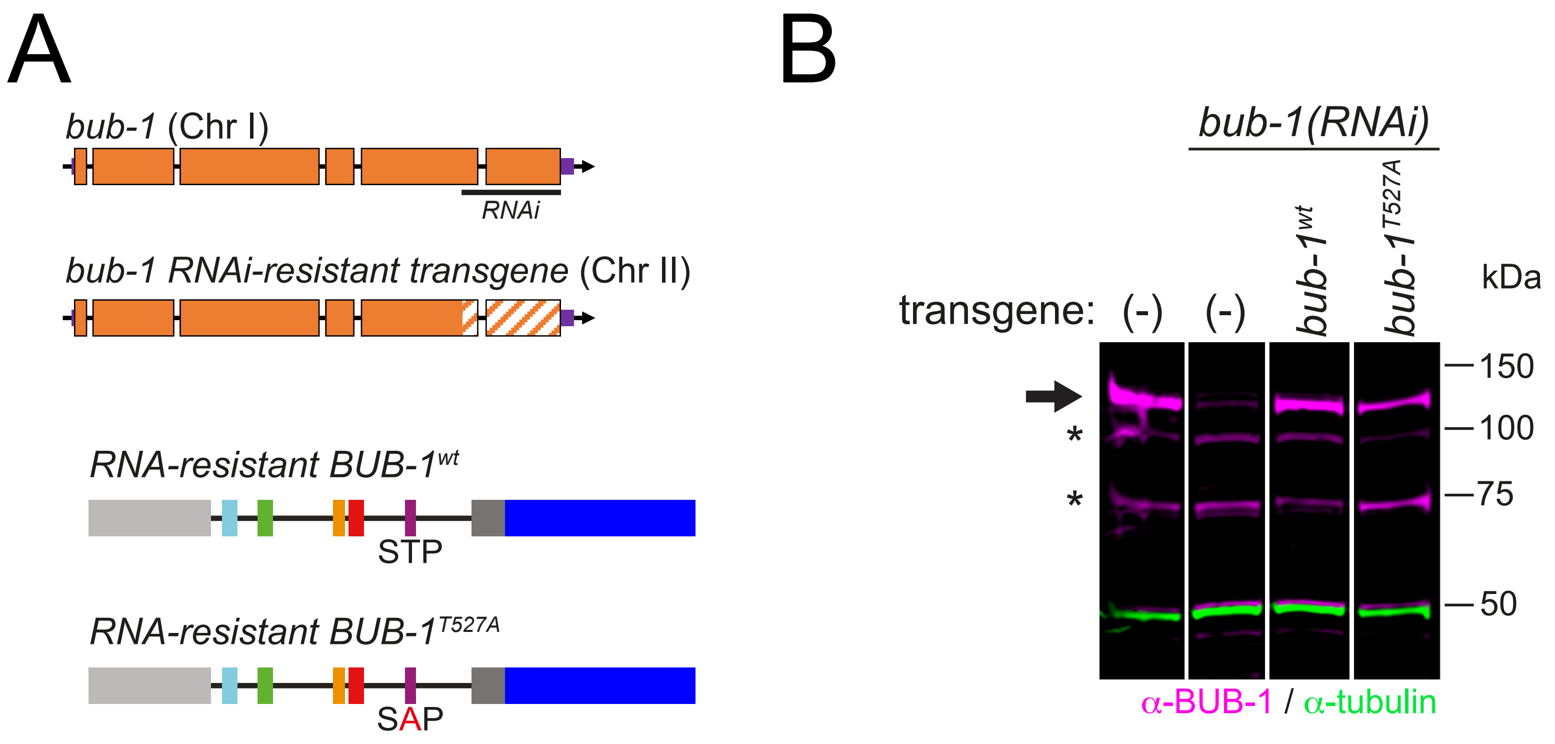

### Figure 9 - Figure supplement 2

#### Figure 9-figure supplement 2

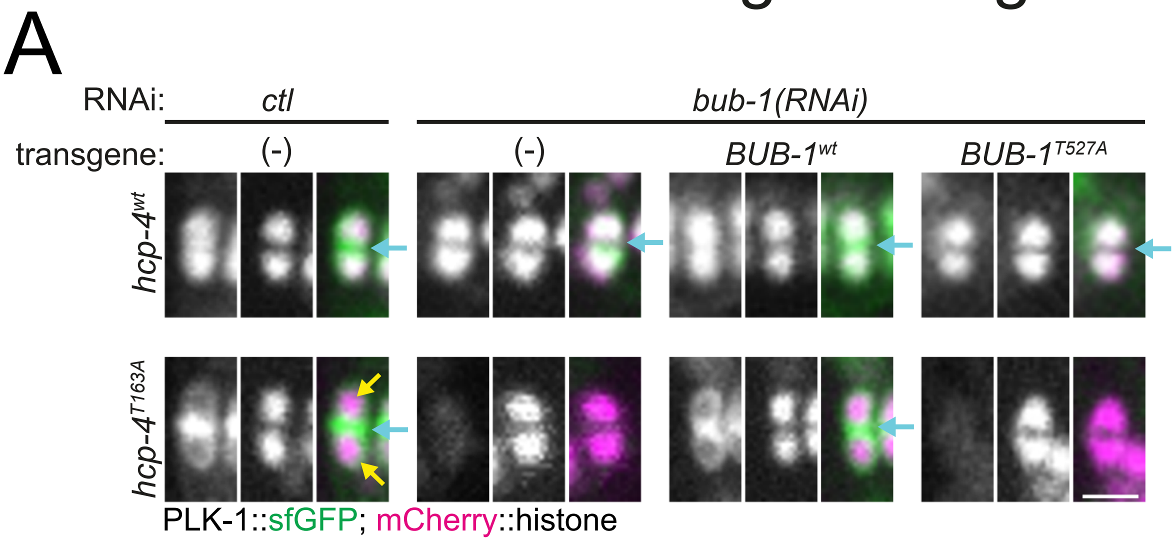


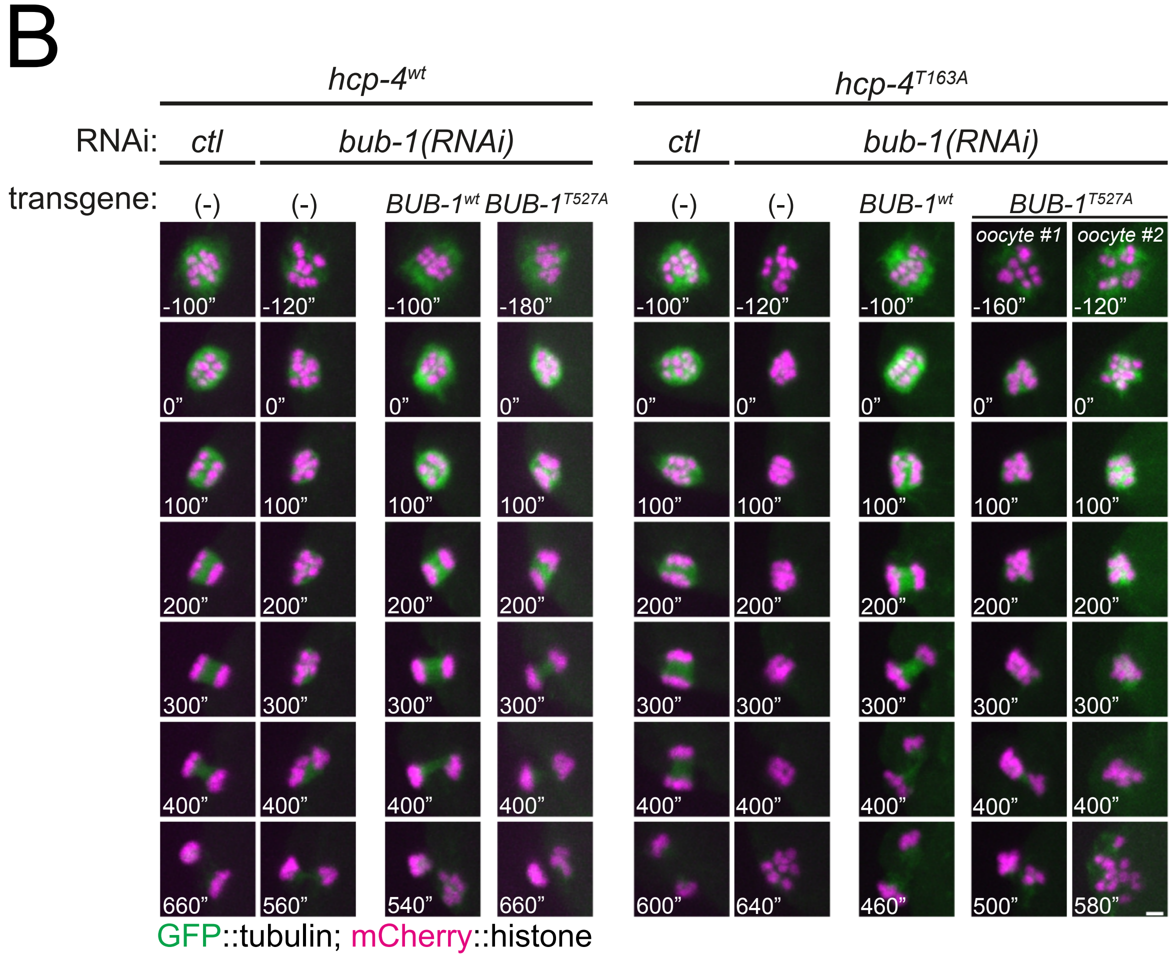
